# Placenta *hIGF1* nanoparticle treatment in guinea pigs mitigates fetal sex dependent FGR-associated effects on kidney structure and blood pressure-related signaling pathways

**DOI:** 10.1101/2025.03.10.642360

**Authors:** Baylea N Davenport, Alyssa Williams, Timothy RH Regnault, Helen N Jones, Rebecca L Wilson

## Abstract

Fetal development in an adverse in utero environment significantly increases the risk of hypertension and cardiovascular disease. The kidneys play a pivotal role in the regulation of blood pressure and cardiovascular function, and perturbations in kidney structure and molecular profile are often demonstrated in offspring born fetal growth restricted (FGR). The aim of this study was to determine whether improving the in utero fetal growth environment with a placental nanoparticle gene therapy would ameliorate FGR-associated dysregulation of fetal kidney development. Using the guinea pig maternal nutrient restriction (MNR) model, we improved placenta efficiency and fetal weight following three placental administrations of a non-viral polymer-based nanoparticle gene therapy from mid-pregnancy (gestational day 35) until gestational day 52. The nanoparticle gene therapy transiently increased expression of *human insulin-like growth factor 1* (*hIGF1*) in placenta trophoblast. Fetal kidney tissue was collected near-term at gestational day 60. Differences in kidney structure, glomeruli size and gene expression of extracellular matrix (ECM) remodeling and blood pressure regulation-related factors were demonstrated in sham-treated FGR fetuses but not observed in FGR fetuses who received placental *hIGF1* nanoparticle treatment. We speculate that mitigating the FGR-associated changes in kidney architecture and molecular profiles might confer protection against increased susceptibility to aberrant kidney physiology in later-life. Overall, this work opens avenues for future research to assess the long-term impact of the placental *hIGF1* nanoparticle gene therapy on cardiovascular function in offspring.

## INTRODUCTION

Fetal growth restriction (FGR), defined as failure of the fetus to meet their genetic growth potential, is diagnosed in 5-10% of pregnancies in the developing world (1). Often FGR is the result of a suboptimal intrauterine environment and developmental adaptations within fetal tissues and organs occur in order for the fetus to survive until birth. These developmental adaptations are collectively known as developmental programming and whilst necessary for in utero survival, can confer long-lasting consequences for an individual’s health. Currently, there are no in utero treatments for FGR. Interventions once FGR has been diagnosed are limited and often include iatrogenic preterm delivery which carries its own inherent health risks. Numerous studies across various human populations indicate that FGR is associated with predisposition to cardiovascular disease, including hypertension and stroke, in adults (2-4). The association between FGR and cardiovascular disease risk in adults is independent of other post-natal life-style factors (5). Studies using various animal models of in utero perturbations that ultimately reduce fetal growth further support the understanding that cardiovascular disease risk is highly influenced by the in utero environment and fetal developmental programming (6-9).

The kidneys play a pivotal role in the regulation of blood pressure and cardiovascular function through an intricate interplay of extrinsic and intrinsic renal mechanisms (10). The regulation process primarily involves the renin-angiotensin-aldosterone system (RAAS) that modulates renal sodium retention and vascular tone (11). When blood pressure is low, the kidneys release renin which catalyzes cleavage of angiotensinogen to angiotensin I. Angiotensin I is then further converted to angiotensin II by angiotensin converting enzyme (ACE). Angiotensin II is a potent vasoconstrictor and through binding with its receptors, raises blood pressure by increasing systemic vascular resistance and stimulating aldosterone secretion, which encourages sodium and water retention. Additionally, the kidney architecture and nephron endowment further contribute to blood pressure homeostasis. Increased fibrotic depositions within the kidneys impair renal blood flow and nephron function, ultimately reducing the ability of the kidneys to regulate blood pressure (12). Lower nephron number is also associated with increased susceptibility to hypertension and chronic kidney disease (13), hypothesized to be due to compensatory mechanisms that alter sodium balance and renal pressure.

Previous studies, predominantly in rodent models, have focused heavily on understanding how FGR compromises kidney architecture, nephron endowment, and gene and protein expression of renal insufficiency markers (6-9). These studies show FGR, as a consequence of various manipulations including uterine vessel ligation, maternal protein or calorie restriction and increase maternal stress, results in reduced glomeruli number (nephron endowment) and increased systemic blood pressure in offspring in later life (6-9). Alterations in kidney morphology and blood pressure are sex dependent with male offspring exhibiting a more predominant phenotype (14). Female offspring only exhibited deficits at a similar magnitude to males when additionally challenged with impaired growth during the lactation period (15). However, kidney development in rodents differs quite significantly from humans. For example, rodent kidneys are morphologically immature at birth (16), making translating knowledge obtained from rodent studies to humans more complex. In humans, kidney development begins early in the first trimester with glomerular function and ultrafiltration beginning around 10-11^th^ week of gestation (17). Nephrogenesis is primarily completed by the 36^th^ week of gestation but continues to mature into the postnatal period (18). Guinea pigs offer the potential to study the developmental origins of deficits to kidney structure and function in a model more reflective of human development (19) as they deliver precocial young whose kidneys have completed development prenatally but continue to mature postnatally (20). Enzymatic activity and hormonal influences on kidney development and function are also similar between guinea pigs and humans (21).

Whilst there are many intrinsic and extrinsic causes of FGR, common to most cases is abnormal placental development and/or function referred to as placental insufficiency (22-24). The placenta is responsible for coordinating the transfer of nutrients, oxygen and waste products between maternal and fetal circulations (25). The transient nature of the placenta makes it an ideal target for FGR treatments as it is discarded after birth. Maternal and fetal circulating levels of growth factors, including Insulin-like 1 Growth Factors (IGF1), are lower in pregnancies complicated by FGR and known to contribute significantly to the pathophysiology of placental insufficiency (26, 27). IGF1 is a master regulator of placental development and function, regulating cell proliferation/ differentiation, vasculature structure, nutrient transport, and hormone production (28, 29). In transgenic mouse models, global deletions in *Igf1* expression results in fetal weight reductions and reduced placental growth (30). We have also shown reduced placental gene expression of *Igf1* in the guinea pig placenta in the maternal nutrient restriction (MNR) model of FGR (31)..

Previous animal-based studies have assessed providing IGF1 protein to the fetus, amniotic fluid or placenta for the treatment of FGR with varied outcomes (32-34). We have demonstrated that repeated delivery of a polymer-based nanoparticle for non-viral, transient *human IGF1* (*hIGF1*) gene delivery to the placenta across the second half of pregnancy improves placenta efficiency (fetal:placental weight ratio), increases fetal blood glucose, reduces fetal blood cortisol and improves fetal weight when compared to sham nanoparticle treated FGR fetuses (31). No off-target nanoparticle expression or plasmid-specific *hIGF1* expression has been observed in fetal tissues indicating that the nanoparticle does not cross the placenta (35, 36). Here, we hypothesize that improving fetal growth with the placenta *hIGF1* nanoparticle gene therapy during critical kidney development windows will mitigate FGR-associated aberrations in kidney structure and function. Our aim was to assess fetal kidney morphology and expression of genes and proteins involved in extracellular matrix remodeling, blood pressure regulation and inflammation/oxidative stress following repeated placenta *hIGF1* nanoparticle gene therapy.

## MATERIALS & METHODS

### Nanoparticle Formation

We have previously published detailed methods of copolymer synthesis and nanoparticle formation (37). Briefly, lyophilized PHPMA_115_-b-PDMEAMA_115_ co-polymer was reconstituted (10 mg/mL) in sterile saline and stored at-20°C. Plasmids were cloned from a pEGFP‐C1 plasmid (Clonetech Laboratories). For the sham nanoparticle, the *CMV* promotor was replaced by a *CYP19A1* promotor and the *GFP* gene was replaced by a non-coding Antisense *GFP* gene. For the *hIGF1* nanoparticle, the *CMV* promotor was replaced by a *CYP19A1* promotor and the *GFP* gene was replaced by a *human IGF1* gene. Nanoparticles were formed by combining 50 µg of plasmid with 75 µL reconstituted polymer and made to a total injection volume of 200 µL with sterile saline under aseptic conditions at room temperature and stored at-20ºC until injection.

### Animal Husbandry and Nanoparticle Delivery

Animal care and usage was approved by the Institutional Animal Care and Usage Committee at the University of Florida (Protocol #202011236). Detailed methods of Animal Husbandry and Nanoparticle Delivery is previously published (31). A schematic of the experimental design is provided in Supplemental Diagram 1. Dams (Dunkin-Hartley) were purchased at 500-550 g (8-9 weeks of age) and allowed to acclimate to the animal facilities for two weeks prior to study initiation. After the acclimatization period, dams were weighed, ranked from heaviest to lightest and systematically assigned to either ad libitum diet (termed Control: n = 6) or maternal nutrient restriction (MNR) diet (n=12) so that the mean weight was no different between the Control and MNR groups. For both Control and MNR dams, water was provided ad libitum. Food (LabDiet diet 5025: 27% protein, 13.5% fat, and 60% carbohydrate as % of energy) intake in the MNR dams was however, restricted to 70% per kilogram body weight of the Control group for 4 weeks prior to mating (dietary conditioning period). Time mating occurred by monitoring the estrus cycle based on changes to the vaginal membrane and detailed in (38). At the end of the dietary conditioning period dams were place with a male guinea pig and changes to the vaginal membrane continued to be monitored daily. Successful mating was presumed to have occurred in the night prior to observing the vaginal membrane as perforated and designated gestational day (GD) 1. Pregnancy checks were performed by ultrasound at approximately GD21 and any dams not pregnant were re-mated. All dams were pregnant within 2 mating attempts. Food restriction (70% per kilogram body weight of the Control group) in the MNR dams continued through to mid‐pregnancy (GD30). From GD30, food was restricted to 90% per kilogram body weight of the Control group to prevent fetal demise and pregnancy loss (39). Control dams were provided food ad libitum.

At GD36±3, dams were anesthetized (Isoflurane: Induction = 3-4% isoflurane with 1-2 min/L oxygen, Maintenance = 1-3% isoflurane with 1-2 min/L oxygen), abdominal hair shaved and skin cleaned using isopropyl alcohol. An ultrasound scan determined the number and location of fetuses. Typically, guinea pigs carry three fetuses, asymmetrically positioned so that one uterine horn contains one fetus, whilst the other uterine horn contains two fetuses. The uterine horn containing one fetus was selected and the ultrasound probe positioned so that neither the fetus nor maternal organs were situated between the maternal body wall and placenta. With the aid of the ultrasound, nanoparticle was delivered by inserting a 30Gx1 needle through the maternal body wall and to the placenta providing a direct intra-placental injection. The needle was removed and placenta monitored briefly for signs of hemorrhaging. Dams assigned to receive sham nanoparticle treatment (n=12; 6 Control dams and 6 MNR dams) were injected with a non-expressing nanoparticle whilst dams assigned to receive the *hIGF1* treatment were injected with a *hIGF1* nanoparticle (n=6 MNR + *hIGF1* dams). Only one placenta per litter was injected. Fetuses whose placentas received the injection were designated the “Directly Treated” fetus. All remaining fetuses in the litter were designated “Indirectly Exposed”. Fetal sex determination at time of first nanoparticle injection was not possible via ultrasound, therefore the sex of the fetus whose placenta received the direct intra-placental injection was random. Intra-placental *hIGF1* or sham nanoparticle injections into the same placenta was repeated at GD 44±3 and GD52±3. Nanoparticle (sham or *hIGF1*) was successfully provided at all 3 timepoints in gestation to all dams. Eight days following the third injection (GD60±3), dams and fetuses were sacrificed by carbon dioxide asphyxiation followed by exsanguination and removal of the heart. Fetuses were sexed based on collection of the gonads (testes for males, ovaries for females). In the dams injected with the *hIGF1* nanoparticle, it was determined by chance that only placentas of male fetuses were directly injected with the *hIGF1* nanoparticle. No placentas of female fetuses were directly injected however, female and other male littermates were indirectly exposed to circulating *hIGF1* nanoparticle. Fetal kidney tissue (Control: n=8 female and n=11 male, MNR: n=5 female and 11 male, MNR + *hIGF1*: n=6 female and 10 male) was weighed and processed for histology (fixed in 4% PFA), RNA extractions (frozen in RNAlater) and flash-frozen in liquid nitrogen. To ensure rigor and reproducibility, all samples were analyzed in a blinded manner.

### Histology and Morphology Analysis

3-5 µm thick paraffin embedded mid-sagittal cross sections of fetal kidney tissue were obtained. Kidney cortex and medulla area was visualized using Masson’s trichrome staining following standard protocols. Collagen deposition was visualized using Sirius Red staining following standard protocols (*Abcam*). Stained sections were imaged on the Axioscan Scanning Microscope (*Zeiss*). Cortex and medulla areas were measured from the Masson’s trichrome stained mid-sagittal cross sections using the Zen Imaging software, v3.9 (*Zeiss*). Quantification of collagen deposition in the cortex and medulla, and glomeruli morphology was determined using a grid-based randomized analysis of the Sirius Red stained sections. Mid-sagittal cross sections were overlayed with a 600 × 900 µm grid in the Zen Imaging software. For collagen deposition, 10 separate grid squares from the cortex and the medulla were randomly selected, imaged and imported into ImageJ Software, v1.54g (40). The Color Deconvolution plugin in ImageJ was used to split images into red, green and blue channels. With the blue channel image selected, percent area was measured using the Threshold and Measure tools. An average percent collagen for the cortex and medulla was then calculated across the 10 images. For glomeruli morphology, all glomeruli were manually counted only in grid sections the overlay the cortex in their entirety. The average number of glomeruli per field was then calculated. In 5 randomly chosen grid fields, all glomeruli without intersection with the grid were manually outlined (range 67-110 glomeruli depending on section), and the mean area (µm^2^) and circularity calculated in the Zen Imaging Software.

### Quantitative PCR (qPCR)

Approximately 30 mg of kidney tissue frozen in RNAlater was homogenized and lysed in RLT‐lysis buffer (*Qiagen*) aided by a tissue homogenizer. RNA was extracted using the RNeasy Mini Plus kit (*Qiagen*) following standard manufacturers protocol. A total of 1 µg of RNA was converted to complementary DNA (cDNA) using the High‐capacity cDNA Reverse Transcription kit (*Applied Biosystems*) and diluted to 1:100. For qPCR, 2.5 µl of cDNA was mixed with 10 µl of PowerUp SYBR green (*Applied Biosystems*), 1.2 µl of primers at a concentration of 10 nM, and water to make up a total reaction volume of 20 µl. Primers for genes of interest specific to the guinea pig genome were purchased commercially (*Millipore* KiCqStart Sybr Primers) or obtained from previously published literature (37, 41, 42). Stability of reference genes *β‐actin, Gapdh* and *Rsp20* (37) in fetal kidney tissue was determined to be 0.011, 0.010 and 0.014, respectively, using Normfinder (43), and gene expression was normalized to the geometric mean of all three. Reactions were performed using the Quant3 Real‐Time PCR System (*Applied Biosystems*), and relative mRNA expression calculated using the comparative CT method with the Design and Analysis Software v2.6.0 (*Applied Biosystems*).

### Western Blots

Fetal kidney tissue was homogenized in ice‐cold RIPA buffer containing protease and phosphatase inhibitors. Protein concentrations determined using Pierce™ Coomassie Plus Assay Kit (*Thermo Fisher Scientific*) following manufacturer’s protocol. 30 μg of protein was mixed with Bolt® SDS Loading Buffer (*Invitrogen*) and Reducing Agent (*Invitrogen*) and denatured by heating at 95°C for 10 min. The lysates and a pre-stain protein ladder (PageRuler, *Thermo Fisher Scientific*) were then run on a 4-12% Tris‐Bis precast gel (*Invitrogen*) following manufacturers protocols and transferred onto nitrocellulose membranes using the Bolt Mini Electrophoresis unit (*Invitrogen*). Membranes were placed into 5% skim‐milk in Tris‐buffered Saline containing Tween 20 (TBS‐T) and incubated overnight at 4°C. Primary antibodies (Oxidative Stress Defense Cocktail 1:1000, *Abcam ab179843*; SOD2: 1:500, *Abcam ab68155*; Deptor: 1:1500, *LSBio C187268*; Total Akt: 1:2000, *Cell Signaling 9272*; Phosphorylated Akt (Ser473): 1:2000, *Cell Signaling 4060*) were applied for 2 h at room temperature or overnight at 4°C, the membranes were then washed 3 times in fresh TBS‐T, and then further incubated with a HRP conjugated secondary (*Cell Signaling 7074* and *7076*, 1:2000) for 2 h at room temperature. Protein bands were visualized by chemiluminescence using SuperSignal West Femto Maximum Sensitivity Substrate (*Thermo Fisher Scientific*) on the Chemidoc Imager (*Bio‐Rad*) and signal intensity of the protein bands calculated using Image Lab software (version 6.1, *Bio-Rad*), normalized to total protein.

### Serum Assays – BUN, Calcium, Creatine and Creatinine

Heparinized fetal blood samples were collected at time of sacrifice by cardiac puncture and plasma obtained by centrifugation. For blood urea nitrogen (BUN), 5 µL of plasma was diluted 1:10 in distilled water analyzed using a colorimetric assay kit (*Invitrogen EIABUN*) following manufacturers protocol. For circulating calcium concentrations, 5 µL of plasma was analyzed using a colorimetric assay kit (*Abcam ab102505*) following manufacturers protocol. For creatine and creatinine concentrations in fetal blood, 10 µL and 15 µL of plasma, respectively was analyzed using a colorimetric assay kit (*BioAssay ECRT-100* and *Arbor Assays KB02*, respectively) following manufacturers protocol. For all assays, samples were run in duplicate, concentrations were calculated from a trendline equation based on standard curve data and optical density corrected by subtracting a blank/background read.

### Statistics

All statistical analyses were performed using SPSS Statistics 29 software. Female and male fetuses were analyzed separately. Distribution assumptions were checked with a Q-Q-Plot. In order to include the possible unmeasurable correlation of the Dam within the litters, statistical significance was determined using generalized estimating equations with gamma as the distribution and log as the link function. Diet and nanoparticle-mediated *hIGF1* treatment were included as main effects, the interaction between diet and nanoparticle-mediated *hIGF1* treatment was included and gestational age and litter size were included as covariates. Direct injection of the placenta compared to indirect exposure to the *hIGF1* or sham nanoparticles was assessed. In the sham treated Control and MNR groups, there was no effect of direct placental injection on any outcomes measured and was therefore removed as a main effect in these groups (31). In MNR + *hIGF1* male fetuses, direct injection of the placenta compared to indirect exposure was significant and therefore remained as a main effect for this group. Statistical significance was considered at P ≤ 0.05. For statistically significant results, a Bonferroni post hoc analysis was performed. Results are reported as estimated marginal means ± 95% confidence interval.

## Results

Maternal, fetal and placental outcomes in these study animals have been reported previously (31). In summary, no adverse maternal health outcomes or pregnancy losses were observed. Average litter size (range: 1-5) did not differ between Control and MNR diets (31). However, one dam (MNR+*hIGF1*) with a litter size of 1 was excluded from the study due to significant physiological differences between monozygotic and polyzygotic pregnancies in guinea pigs (44). No resorptions were recorded (31). There was no difference in the number of female or male fetuses between Control or MNR diet (31). However, it was determined at time of sacrifice that only placentas of male fetuses had received a direct placental injection of *hIGF1* nanoparticle. Repeated placental *hIGF1* nanoparticle treatment resulted in gene expression of plasmid-specific *hIGF1* in all placentas of the litter; directly injected placentas from male fetuses had higher levels of *hIGF1* than indirectly exposed female and male littermates (31). Plasmid-specific *hIGF1* expression was not detected in the sham nanoparticle treated Control or MNR placentas. Near-term fetal weight was lower in the sham treated MNR dams when compared to sham treated Control dams (31). However, near-term fetal weight in the *hIGF1* treated MNR dams was comparable to sham treated Control dams (31).

### MNR resulted in changes to the fetal kidney architecture which was prevented by placental

#### *hIGF1* nanoparticle treatment depending on fetal sex

In female fetuses, neither absolute kidney weight (both kidneys combined) nor kidney weight corrected for fetal weight was different between sham Control, sham MNR and indirectly exposed MNR+*hIGF1* (Supplemental Figure S1A and B). In male fetuses, absolute kidney weight (both kidneys combined) was reduced in the sham MNR when compared to sham Control and MNR+*hIGF1* (Supplemental Figure 1C). However, once corrected for fetal weight, there was no difference in kidney weight between sham Control, sham MNR and MNR+*hIGF1* (Supplemental Figure S1D). Mid-sagittal cortex area was not different with MNR or placental *hIGF1* nanoparticle treatment in female and male fetuses (Figure 1A and 1B, respectively). Mid-sagittal medulla area was lower in sham MNR female and male fetuses when compared to sham Control (Figure 1C and 1D, respectively). Reduced mid-sagittal medulla area resulted in increased cortex:medulla area ratios in the sham MNR fetuses compared to sham Control for both fetal sex (Figure 1E and 1F). In MNR+*hIGF1* female fetuses whose placentas were indirectly exposed to *hIGF1* nanoparticle, mid-sagittal medulla area was reduced compared to sham Control and similar to sham MNR (Figure 1C). However, in MNR+*hIGF1* male fetuses whose placentas were both indirectly exposed and directly injected with *hIGF1* nanoparticle, mid-sagittal medulla area was increased compared to sham MNR, and comparable to sham Control (Figure 1D).

**Figure 1.**
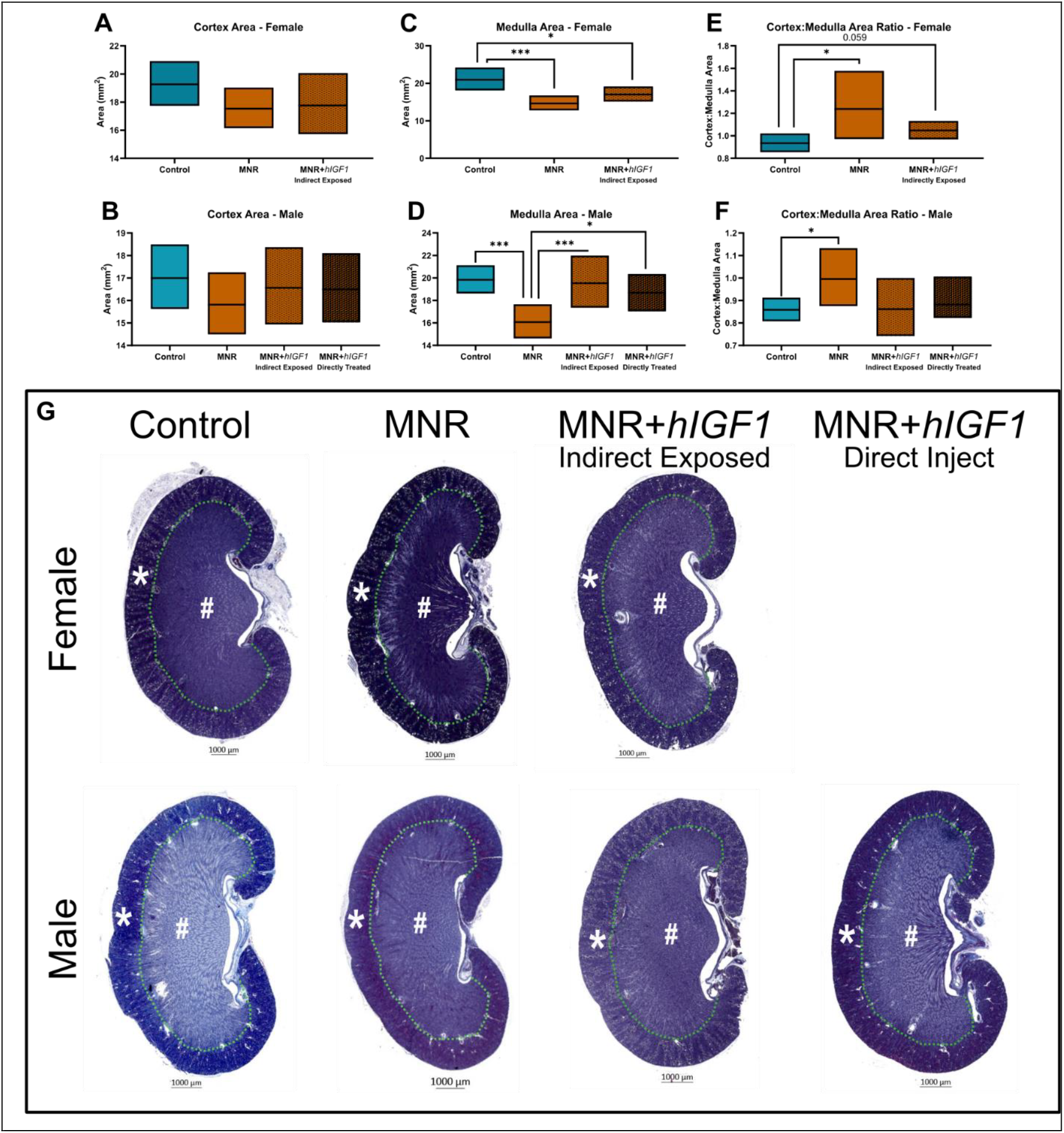
Effects of maternal nutrient restriction (MNR) and repeated placental *hIGF1* nanoparticle treatment (MNR+*hIGF1*) on fetal kidney morphology. **A**. In female fetuses, mid-sagittal cortex area was not different with MNR or placental *hIGF1* nanoparticle treatment. **B**. In male fetuses, mid-sagittal cortex area was not different with MNR or placental *hIGF1* nanoparticle treatment. **C**. In female fetuses, mid-sagittal medulla area was lower in sham MNR compared to sham Control. In MNR+*hIGF1* female fetuses whose placentas were indirectly exposed, mid-sagittal medulla area was reduced compared to sham Control and similar to sham MNR. **D**. In male fetuses, mid-sagittal medulla area was lower in sham MNR compared to sham Control. In MNR+*hIGF1* male fetuses mid-sagittal medulla area was increased compared to sham MNR, and comparable to sham Control. **E**. Cortex:medulla area ratio was increased in sham MNR female fetuses compared to sham Control. **F** Cortex:medulla area ratio was increased in sham MNR male fetuses compared to sham Control. **G**. Representative mid-sagittal cross sections of the fetal kidney stained with Masson’s Trichrome; Cortex (star) and Medulla (asterix) are delineated by a green dashed line. Control: n = 6 dams (8 female and 11 male fetuses), MNR: n = 6 dams (5 female and 11 male fetuses), MNR+*hIGF1*: n = 5 dams (6 female and 10 male fetuses). Data are estimated marginal means ± 95% confidence interval. *P≤0.05; **P≤0.01. ***P≤0.001

Nephron number (number of glomeruli) was not different in either female or male fetuses with MNR or placental *hIGF1* nanoparticle treatment (Figure 2A and 2B, respectively). Mean glomeruli area of female fetuses was similar between sham Control, sham MNR and MNR+*hIGF1* (Figure 2C). In male fetuses, mean glomeruli area was lower in sham MNR compared to sham Control (Figure 2D). In both indirectly exposed and direct treated MNR+*hIGF1* males, mean glomeruli area was similar to both sham Control and sham MNR (Figure 2D). Glomeruli circularity was increased in indirectly exposed MNR+*hIGF1* female fetuses and directly treated MNR+*hIGF1* male fetuses compared to sham Control (Figure 2E and 2F, respectively).

**Figure 2.**
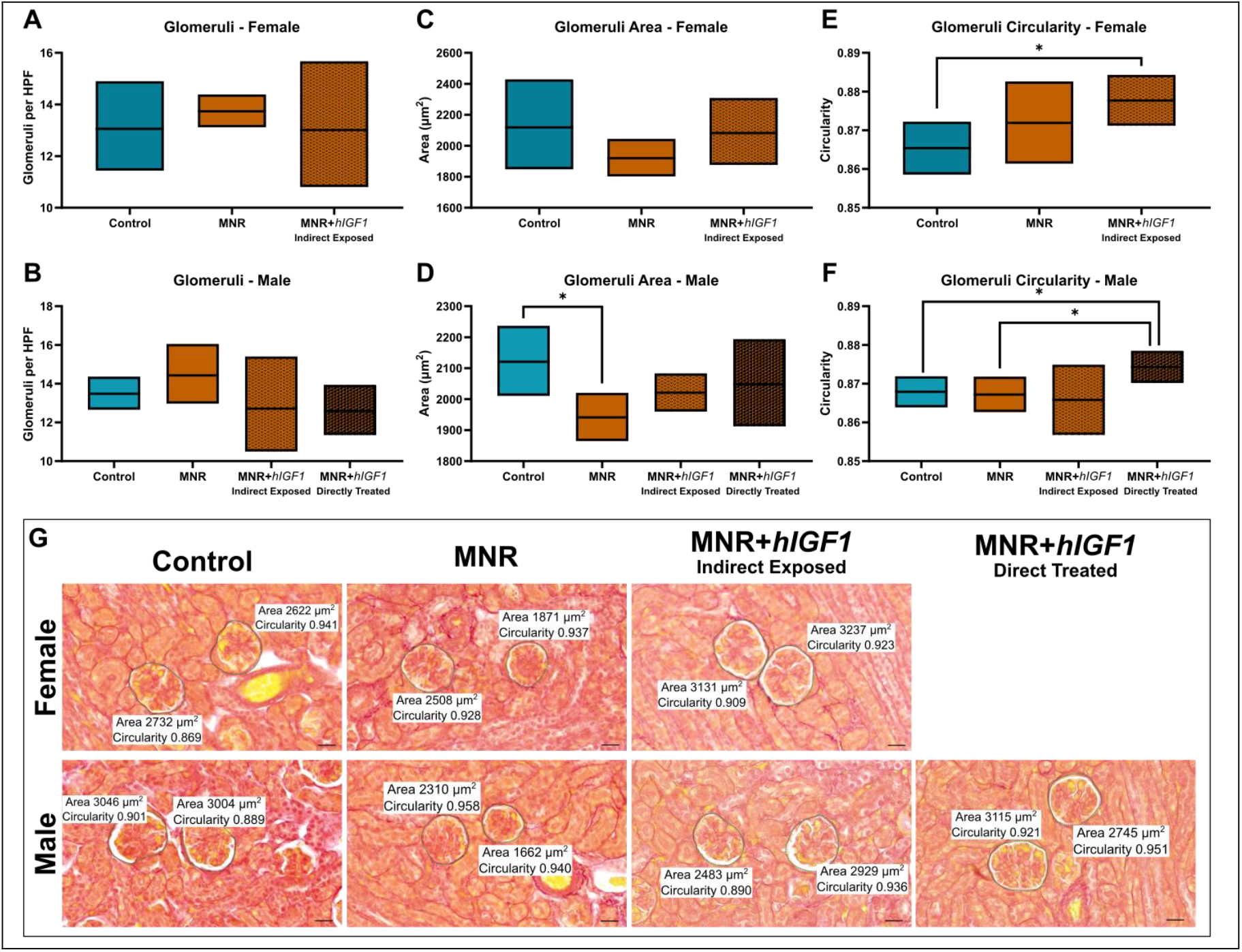
Effects of maternal nutrient restriction (MNR) and repeated placental *hIGF1* nanoparticle treatment (MNR+*hIGF1*) on fetal kidney glomeruli morphology. **A**. In female fetuses, nephron number (number of glomeruli) was not different with MNR or placental *hIGF1* nanoparticle treatment. **B**. In male fetuses, nephron number was not different with MNR or placental *hIGF1* nanoparticle treatment. **C**. Mean glomeruli area of female fetuses was similar between sham Control, sham MNR and MNR+*hIGF1*. **D**. In male fetuses, mean glomeruli area was lower in sham MNR compared to sham Control. In both indirectly exposed and direct treated MNR+*hIGF1* male fetuses, mean glomeruli area was similar to both sham Control and sham MNR. **E**. In female fetuses, glomeruli circularity was increased in indirectly exposed MNR+*hIGF1* compared to sham Control. **F**. In male fetuses, glomeruli circularity was increased in directly treated MNR+*hIGF1* fetuses compared to sham Control, sham MNR and indirectly exposed MNR+*hIGF1*. **G**. Representative images of picrosirius red stained glomeruli (circled with a green dashed line) assessed for area and circularity. Scale bar = 50 µm. Control: n = 6 dams (8 female and 11 male fetuses), MNR: n = 6 dams (5 female and 11 male fetuses), MNR+*hIGF1*: n = 5 dams (6 female and 10 male fetuses). Data are estimated marginal means ± 95% confidence interval. *P≤0.05.

### In female but not male fetuses, MNR increased kidney gene expression of ECM remodeling factors which was prevented by placental *hIGF1* nanoparticle treatment

Increased production of ECM proteins and collagen in the kidneys can interfere with the filtering capacity of the glomeruli and increases glomerulosclerosis (45). Increased collagen deposition around the glomeruli and tubules was not observed with either MNR or placental *hIGF1* nanoparticle treatment (Supplemental Figure S2 and S3, respectively). In female fetuses, renal mRNA expression of *Transforming Growth Factor beta* (*Tgfb*), *Matrix Metalloproteinase 2* (*Mmp2*) and *TIMP metalloproteinase inhibitor 1* (*Timp1*) was increased in sham MNR fetuses compared to sham Control (Figure 3A-C, respectively). mRNA expression of *Tgfb* and *Mmp2* were decreased in the indirectly exposed MNR+*hIGF1* female fetuses compared to sham MNR, and comparable to levels in sham Control (Figure 3A and B, respectively). Renal expression of *Timp2* was decreased in the indirectly exposed MNR+*hIGF1* female fetuses compared to sham Control whilst expression of *Connective Tissue Growth Factor* (*Ctgf*) was increased in the indirectly exposed MNR+*hIGF1* female fetuses compared to sham Control and sham MNR (Figure 3D and E, respectively). In male fetuses, renal mRNA expression of *Tgfb, Mmp2, Timp1* and *Ctgf* was similar between the groups (Figure 3F-H and J, respectively). Compared to sham Control, renal mRNA expression of *Timp2* was decreased in the sham MNR, and indirectly exposed and directly treated MNR+*hIGF1* male fetuses (Figure 3I).

**Figure 3.**
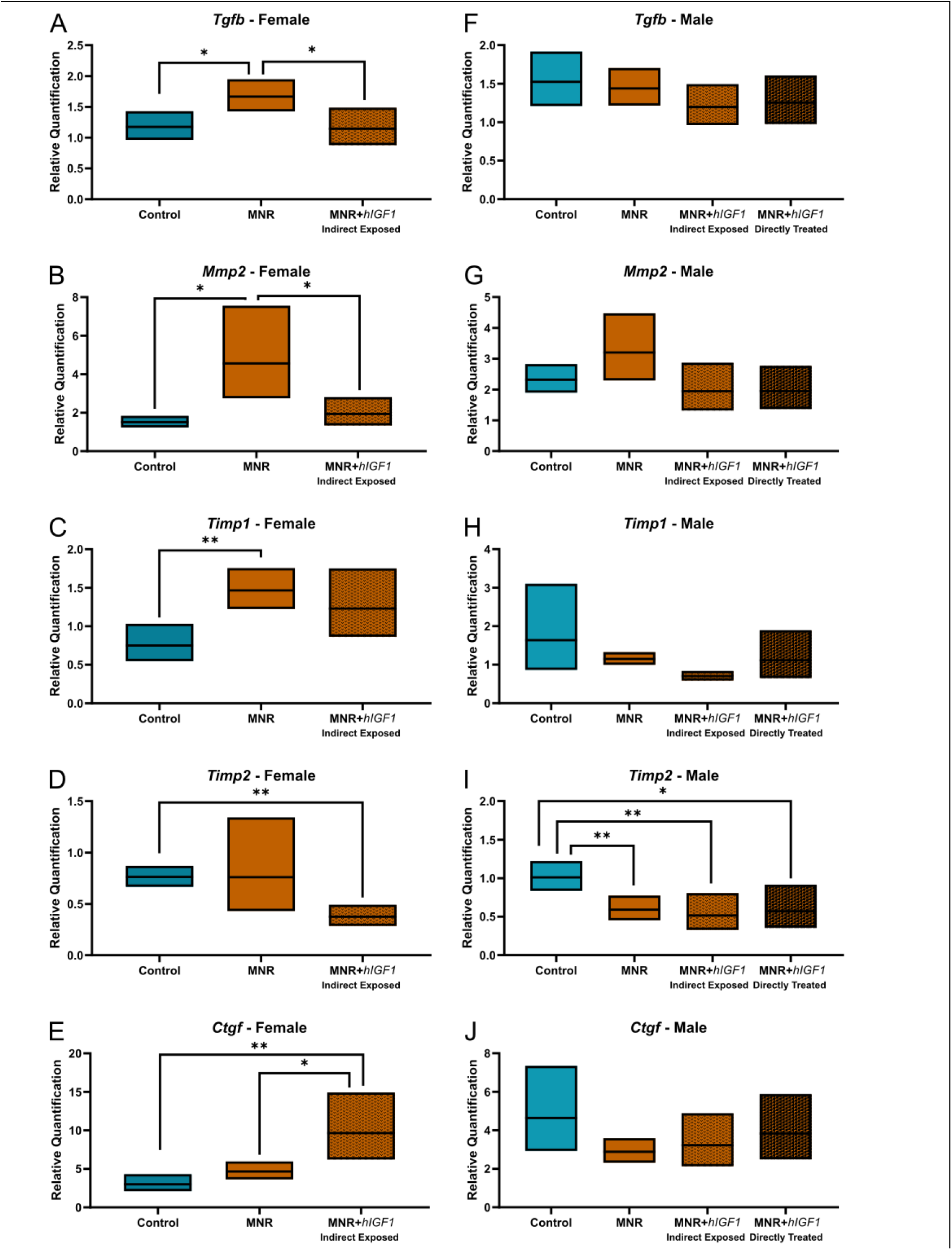
Effects of maternal nutrient restriction (MNR) and repeated placental *hIGF1* nanoparticle treatment (MNR+*hIGF1*) on renal mRNA expression of genes involved in extracellular matrix remodeling. **A**. In female fetuses, renal mRNA expression of *Transforming Growth Factor beta* (*Tgfb*) was increased in sham MNR fetuses compared to sham Control and decreased in the MNR+*hIGF1* fetuses compared to sham MNR. **B**. In female fetuses, renal mRNA expression of *Matrix Metalloproteinase 2* (*Mmp2*) was increased in sham MNR fetuses compared to sham Control and decreased in the MNR+*hIGF1* fetuses compared to sham MNR. **C**. In female fetuses, renal mRNA expression of *TIMP metalloproteinase inhibitor 1* (*Timp1*) was increased in sham MNR fetuses compared to sham Control and decreased in the MNR+*hIGF1* fetuses compared to sham MNR. **D**. Renal expression of *Timp2* was decreased in the MNR+*hIGF1* female fetuses compared to sham Control. **E**. Renal expression of *Connective Tissue Growth Factor* (*Ctgf*) was increased in the MNR+*hIGF1* female fetuses compared to sham Control and sham MNR. **F**. In male fetuses, renal mRNA expression of *Tgfb* was similar across the groups. **G**. In male fetuses, renal mRNA expression of *Mmp2* was similar across the groups. **H**. In male fetuses, renal mRNA expression of *Timp1* was similar across the groups. **I**. Compared to sham Control, renal mRNA expression of *Timp2* was decreased in the sham MNR and MNR+*hIGF1* nanoparticle male fetuses. **J**. In male fetuses, renal mRNA expression of *Ctgf* was similar across the groups. Control: n = 6 dams (8 female and 11 male fetuses), MNR: n = 6 dams (5 female and 11 male fetuses), MNR+*hIGF1*: n = 5 dams (6 female and 10 male fetuses). Data are estimated marginal means ± 95% confidence interval. *P≤0.05; **P≤0.01.

### MNR resulted in fetal sex-dependent alterations in kidney RAS gene expression

Alterations in the renal renin-angiotensin system have been shown in rat models of FGR with changes correlating with increased blood pressure and altered renal development in offspring (46-48). In female fetuses, renal mRNA expression of *Angiotensinogen* (*Agt*) was increased in the sham MNR and indirectly exposed MNR+*hIGF1* fetuses compared to sham Control (Figure 4A). Renal mRNA expression of *Renin* (*Ren*) was increased in the indirectly exposed MNR+*hIGF1* female fetuses compared to the sham Control and sham MNR fetuses (Figure 4B). mRNA expression of *Angiotensin Converting Enzyme* (*Ace*) trended towards a decrease in the sham MNR female fetuses when compared to sham Control (Figure 4C). *Ace* expression was increased in the indirectly exposed MNR+*hIGF1* female fetuses when compared to sham MNR resulting in expression comparable to sham Control (Figure 4C). Expression of *Renin Binding Protein* (*RenBP*) and *Angiotensin Type 1 Receptor* (*AgtR1*) was similar in female fetuses across the three groups (Supplemental Table S1). In male fetuses, renal mRNA expression of *Agt* was similar across the groups (Figure 4D). Renal mRNA expression of *Ren* was increased in the directly treated MNR+*hIGF1* male fetuses compared to sham MNR (Figure 4E). mRNA expression of *Ace* was decreased in sham MNR compared to sham Control, and increased in the directly treated MNR+*hIGF1* male fetuses when compared to sham MNR (Figure 4F). Expression of *RenBP* was similar across the groups, whilst expression of *AgtR1* was decreased in the directly treated MNR+*hIGF1* male fetuses compared to sham MNR (Supplemental Table S1). Angiotensin *Type 2 Receptor* (*AgtR2*) was not expressed (no detectable expression after 40 qPCR cycles) in either female or male fetal kidneys (data not shown).

**Figure 4.**
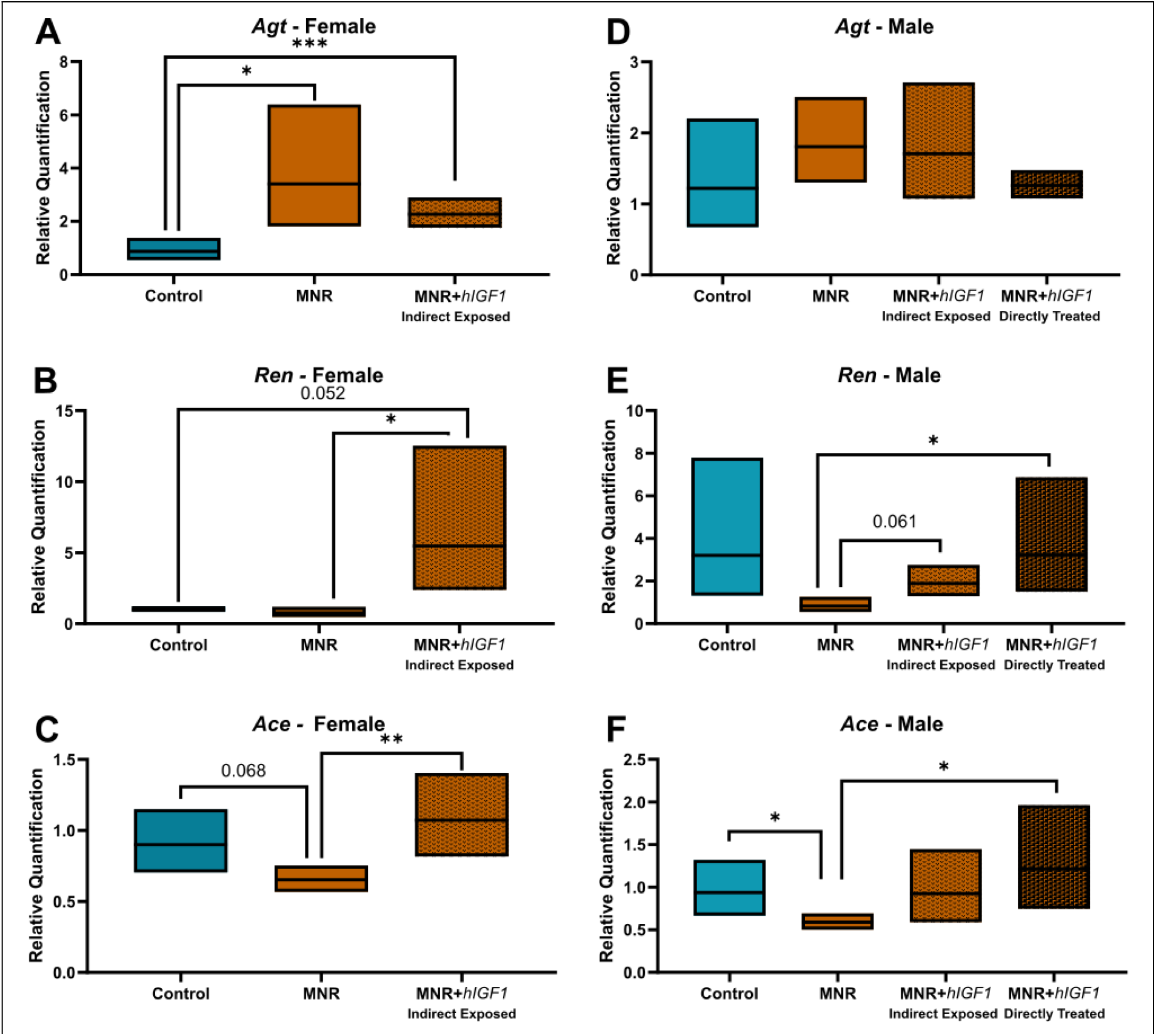
Effects of maternal nutrient restriction (MNR) and repeated placental *hIGF1* nanoparticle treatment (MNR+*hIGF1*) on renal mRNA expression of Renin-Angiotensin System genes. **A**. In female fetuses, renal mRNA expression of *Angiotensinogen* (*Agt*) was increased in the sham MNR and MNR+*hIGF1* fetuses compared to sham Control. **B**. Renal mRNA expression of *Renin* (*Ren*) was increased in the MNR+*hIGF1* female fetuses compared to sham Control and sham MNR fetuses. **C**. mRNA expression of *Angiotensin Converting Enzyme* (*Ace*) trended towards a decrease in the sham MNR female fetuses when compared to sham Control. *Ace* expression was increased in the MNR+*hIGF1* female fetuses when compared to sham MNR. **D**. In male fetuses, renal mRNA expression of *Agt* was similar across the groups. **E**. Renal mRNA expression of *Ren* was increased in the MNR+*hIGF1* males compared to sham MNR males, particularly in those whose placentas were directly treated. **F**. mRNA expression of *Ace* was decreased in sham MNR male fetuses compared to sham Control. *Ace* expression was increased in the directly treated MNR+*hIGF1* male fetuses compared to sham MNR. Control: n = 6 dams (8 female and 11 male fetuses), MNR: n = 6 dams (5 female and 11 male fetuses), MNR+*hIGF1*: n = 5 dams (6 female and 10 male fetuses). Data are estimated marginal means ± 95% confidence interval. *P≤0.05; **P≤0.01. ***P≤0.001

### No indicators of increased inflammation or oxidative stress were observed in the near-term fetal kidneys

Increased inflammation and oxidative stress as well as antioxidant loss has been associated with kidney disease progression and cardiovascular disease in humans (49). In female guinea pig fetuses, renal mRNA expression of *Interleukin 6* (*Il6*) and its receptor: *Il6R* were similar between the groups (Supplemental Table S1). mRNA expression of *Hypoxia Inducible Factor 1a* (*Hif1a*) was increased in sham MNR and indirectly exposed MNR+*hIGF1* female fetuses compared to sham Control (Supplemental Table S1). Expression of *Heat Shock Protein Family Member A 5* (*Hspa5*) was increased in the sham MNR female fetuses compared to sham Control (Supplemental Table S1). *Hspa5* was decreased in the indirectly exposed MNR+*hIGF1* female fetuses compared to sham MNR and comparable to levels in sham Control (Supplemental Table S1). Renal protein expression of Thioredoxin was reduced in the indirectly exposed MNR+*hIGF1* female fetuses compared to sham Control and sham MNR (Supplemental Table S1). Protein expression of Catalase and Superoxide Dismutase (SOD2) was similar across the three groups (Supplemental Table S1). In male fetuses, renal mRNA expression of *Il6, Il6R, Hif1a* and *Hspa5* was comparable between sham Control, sham MNR and MNR+*hIGF1* fetuses. Renal protein expression of Thioredoxin and Catalase was reduced in the indirectly exposed and directly treated MNR+*hIGF1* male fetuses compared to sham Control and sham MNR (Supplemental Table S1). Protein expression of SOD2 was reduced in the sham MNR and indirectly exposed MNR+*hIGF1* nanoparticle male fetuses compared to sham Control (Supplemental Table 1). SOD1 protein was not detected in kidney tissue from female and male fetuses (data not shown).

### MNR increased circulating blood urea nitrogen (BUN) in female fetuses and decreased circulating creatinine in male fetuses, but not with placental *hIGF1* nanoparticle treatment

BUN, calcium, creatine and creatinine were measured in fetal plasma to indirectly assess fetal kidney function (50). In female fetuses, plasma BUN was increased in sham MNR compared to sham Control (Figure 5A). BUN was reduced in the indirectly exposed MNR+*hIGF1* female fetuses compared to sham MNR (Figure 5A). Plasma creatine was increased in the sham MNR and indirectly exposed MNR+*hIGF1* female fetuses when compared to sham Control (Figure 5B). There was no difference in circulating creatinine or calcium in female fetuses between any of the groups (Figure 5C and 5D, respectively). In male fetuses, plasma concentrations of BUN were comparable across the groups (Figure 5E). Plasma creatine was increased in the sham MNR, and indirectly exposed and direct treated MNR+*hIGF1* male fetuses compared to sham Control (Figure 5F). Plasma creatinine was reduced in the sham MNR male fetuses compared to sham Control (Figure 5G). Creatinine was increased in the directly treated MNR+*hIGF1* male fetuses compared to sham MNR and at similar levels as sham Control (Figure 5G). Plasma concentrations of calcium in male fetuses were similar across the groups (Figure 5H).

**Figure 5.**
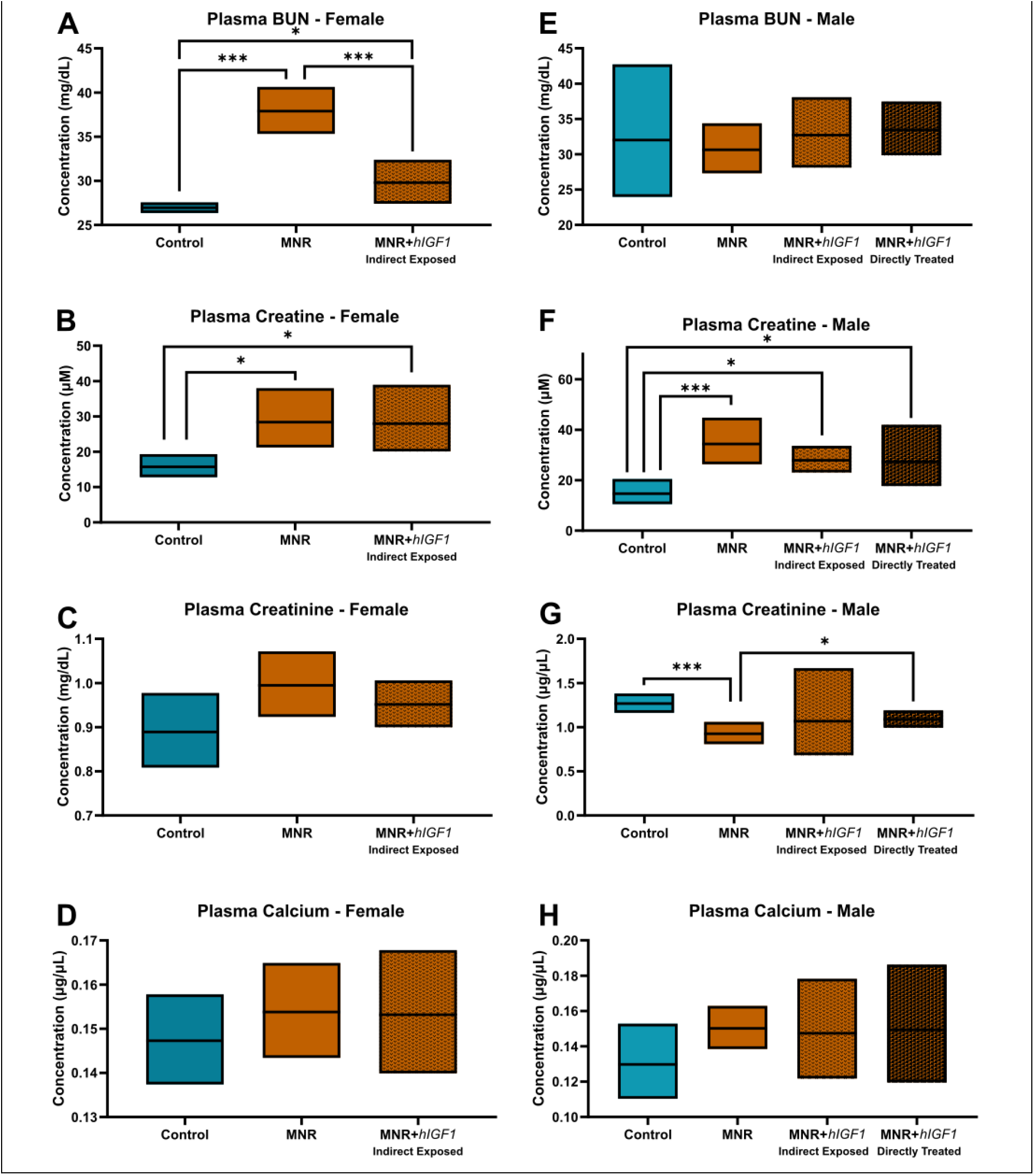
Effects of maternal nutrient restriction (MNR) and repeated placental *hIGF1* nanoparticle treatment (MNR+*hIGF1*) on circulating markers of kidney function. **A**. In female fetuses, plasma Blood Urea Nitrogen (BUN) was increased in sham MNR compared to sham Control but reduced back to comparable levels to Control in the indirectly exposed MNR+*hIGF1* female fetuses. **B**. Plasma creatine was increased in the sham MNR and MNR+*hIGF1* female fetuses when compared to sham Control. **C**. In female fetuses there was no difference in circulating creatinine between any of the groups. **D**. In female fetuses there was no difference in circulating calcium between any of the groups. **E**. In male fetuses, plasma concentrations of BUN were comparable across the groups. **F**. Plasma creatine was increased in the sham MNR and MNR+*hIGF1* male fetuses compared to sham Control. **G**. Plasma creatinine was reduced in the sham MNR fetuses compared to sham Control but increased in the directly treated MNR+*hIGF1* male fetuses. **H**. In male fetuses, plasma concentrations of calcium were comparable across the groups. Control: n = 6 dams (8 female and 11 male fetuses), MNR: n = 6 dams (5 female and 11 male fetuses), MNR+*hIGF1*: n = 5 dams (6 female and 10 male fetuses). Data are estimated marginal means ± 95% confidence interval. *P≤0.05; **P≤0.01. ***P≤0.001

## Discussion

In our previous placental *hIGF1* nanoparticle studies (31, 37, 51, 52) that utilize the well characterized guinea pig maternal nutrient restriction (MNR) model of FGR we demonstrated the ability to positively influence fetal growth by enhancing placenta efficiency (31). Here, we furthered this knowledge and assessed the impact of our intervention on fetal kidney development and expression of genes and proteins involved in extracellular matrix (ECM) remodeling, blood pressure regulation and oxidative stress. We demonstrated that FGR was associated with decreased mid-sagittal medullar area, reduced glomerular size and fetal sex-specific changes to ECM remodeling and RAS gene expression. Kidney gene expression profiles and morphology in the FGR fetuses whose placentas were treated with the *hIGF1* nanoparticle were more similar to Control indicating the ability to indirectly impact kidney development through improving the in utero environment. We speculate that preventing FGR-associated changes in kidney RAS and ECM remodeling pathway expression might confer protection against increased susceptibility to abnormal blood pressure regulation and cardiovascular disease in later life. Overall, the findings illuminate the transformative potential of placental-specific *hIGF1* nanoparticle gene therapy in mitigating the impacts of placenta insufficiency on fetal kidney development and addressing FGR-related cardiovascular programming challenges.

Increased risk of developing hypertension and cardiovascular disease in later life is strongly associated with deficits in kidney morphology, including size of the medulla (53, 54). In the sham treated FGR fetuses, medulla area was reduced when compared to sham treated Control fetuses. The medulla plays a crucial role in concentrating urine and maintaining fluid and electrolyte balance (55). A diminished medullary area may have implications on kidney function and lead to impaired ability to regulate sodium excretion and water reabsorption, and consequently disrupt blood pressure control. In the male FGR fetuses whose placentas were treated with the *hIGF1* nanoparticle, medullar area was restored to size comparable to the sham Control males. In rats, continuous exposure of the kidney to a synthetic glucocorticoid receptor agonist from birth to postnatal day 11 selectively impairs development of renal outer medulla (53). We have previously shown in these male fetuses, increased blood cortisol concentrations in the sham MNR group which were reduced to Control levels in the MNR+*hIGF1* group (31) supporting a possible mechanistic link between kidney medulla development, circulating cortisol and placental *hIGF1* intervention worth further exploration. Increased medulla area in the female fetuses that were indirectly exposed to *hIGF1* nanoparticle was not observed despite a trend for reduced circulating cortisol in these female fetuses (31). However, this likely reflects a dosage response which is currently under investigation with direct treatment of female placentas.

Reduced nephron endowment following placental insufficiency resulting in FGR has also been suggested to be an underlying cause in the development of essential hypertension. In animal studies, fewer glomeruli are often characterized in FGR offspring at various post-natal time points (9, 14, 15, 56). Similarly, in humans, birth weight, glomerular number and blood pressure are strongly associated (57, 58). Reduced glomeruli number has been characterized in the FGR guinea pig at 8-weeks of age (9). We did not observe a difference in the number of glomeruli in the near-term fetal guinea pig with either MNR or MNR+*hIGF1* nanoparticle treatment suggesting nephron deficits may not be established before birth but possibly during postnatal life. Glomeruli size was reduced, particularly in the sham treated FGR male fetuses, indicating less surface area for filtration and the potential for reduced glomerular filtration rate (GFR). Overall these findings suggest that reduced glomerular size in combination with a potential postnatal loss of nephrons, may impair filtration capacity and contribute to long-lasting disruptions in blood pressure regulation that are characteristic of FGR guinea pig models in adulthood (9, 59).

In addition to glomeruli size and number, increased production of ECM proteins and collagen in the kidneys can interfere with the filtering capacity of the glomeruli, which in turn decreases filtration and increases glomerulosclerosis (45). Our results did not demonstrate significant difference in collagen deposition in the cortex or medulla with either MNR or MNR+*hIGF1* nanoparticle treatment. However, this is likely reflective of developmental stage as excessive ECM deposition from birth would likely confer very poor survival of the offspring. In female fetuses, we did identify increased mRNA expression of ECM remodeling factors in the MNR group. Increased mRNA expression of ECM remodeling factors in the kidneys of MNR male fetuses was not observed, an outcome similar to previous studies in rats which have shown molecular changes in ECM remodeling pathways in FGR female pups but not FGR male pups (14, 15). Changes in expression of *Tgfb, MMP*s and *TIMP*s have been shown to precede the development of glomerulosclerosis and interstitial fibrosis following renal damage (60, 61). Furthermore, increased collagen deposition in the kidneys in 8-week-old guinea pigs following FGR has previously been shown (9), although fetal sex as a biological variable was not included. Whilst our results show there was no marked fibrosis or glomerulosclerosis in the fetal kidneys, there is a molecular vulnerability beginning in utero that may contribute to increased susceptibility to glomerular damage in the future. Indirect exposure of the placenta to the *hIGF1* nanoparticle in the female fetuses resulted in kidney expression of ECM remodeling factors that were more similar to Control further indicating the ability to indirectly influence fetal kidney gene expression through modification of placental development/function.

Placental treatment with the *hIGF1* nanoparticle also indirectly modified expression of RAS genes and oxidative stress proteins in the fetal kidneys. Angiotensin II (Ang II) is a potent vasoconstrictor, and imbalances in the production of Ang II lead to heightened vasoconstriction, sodium retention and increased blood pressure (62). Similarly, increased oxidative stress in the kidneys not only impairs renal function by damaging nephrons, but also enhances RAS activity (49). Hence, therapeutic strategies for managing hypertension often focus on targeting RAS overactivity and/or oxidative stress (63). In our study, both FGR and placental *hIGF1* nanoparticle treatment modified kidney gene expression of *Agt, Ren* and *Ace*, depending on fetal sex. In particular, expression of *Ace* was lower in sham treated FGR male fetuses and trended towards being lower in sham treated FGR female fetuses compared to Control suggesting a shift in the regulation of RAS in the kidneys. Typically, increased *Ace* expression and/or activity is associated with hypertension, thus we expected to see higher levels in the FGR fetuses. Additionally, we expected to see more pronounced differences in expression of oxidative stress proteins between sham treated FGR and Control fetuses. Conclusions as to how these molecular changes in RAS and oxidative stress protein expression are impacting kidney function are limited because urine could not be collected at the fetal timepoint. Surrogate markers of kidney function in fetal blood were measured including BUN and creatinine. However, circulating blood factors are also heavily influenced by non-renal causes, such as liver function and body composition (64-66). Thus, further studies in offspring whose placentas were treated with the *hIGF1* nanoparticle, and from which urine can be collected, are underway to more comprehensively assess kidney function and to fully elucidate the implications of indirectly modifying kidney gene and protein expression.

The influence of fetal sex on pregnancy outcomes and subsequent post-natal health is well established (67-69). In pregnancy, male fetuses are at higher risk of poorer outcomes than female fetuses (67). Postnatally males exhibit more severe cardiovascular pathologies (68, 69) and is supported by animal studies which often show increased blood pressure in male offspring but not female offspring following FGR (14, 15, 59, 70, 71). Sexually dimorphic responses in blood pressure are likely driven by sex hormones as gonadectomy has confirmed the ability of testosterone removal to reduce blood pressure whilst estrogen removal increases blood pressure (72, 73). In guinea pigs specifically, male fetuses have the capacity to biosynthesize androgens between day 24 and 30 of gestation (74). Additionally, circulating testosterone and DHT are higher in male fetuses compared to female fetuses from gestational day 35 (75). Therefore, it is highly plausible that fetal sex hormones have some influence on the outcomes we observed. One limitation to further explore this hypothesis, however, is the inability to study the impact of the *hIGF1* nanoparticle gene therapy in female fetuses whose placentas were directly injected. As most human pregnancies are singleton with one placenta, one fetus in the litter was chosen to receive a direct sham or *hIGF1* nanoparticle injection to study effects of the full direct treatment. The remaining fetuses in the litter were used to investigate possible circulating indirect exposure. However, the sex of the fetus which received the direct injection cannot be controlled nor reliably determined at mid-pregnancy via ultrasound. Further studies in non-human primates are being performed to address translational aspects of the *hIGF1* nanoparticle gene therapy including dose, which are difficult to ascertain in litter bearing species (76).

In conclusion, we investigated the effects of FGR and a placental-specific *hIGF1* nanoparticle treatment on fetal kidney development. The focus was on kidney structure and gene and protein expression of factors involved in ECM remodeling, blood pressure regulation and oxidative stress. Our findings further corroborate previous investigations supporting the in uteroin utero origins of vulnerabilities in the kidneys that contribute to hypertension and cardiovascular disease risk in later life. Additionally, we have further evidence to support the beneficial impact of our placental *hIGF1* nanoparticle gene therapy on fetal growth and development. The differences in kidney development and molecular programming between female and male fetuses further emphasizes the need for sex-specific approaches in understanding and treating FGR-induced cardiovascular disturbances. Overall, our work opens avenues for future research to assess the long-term impact of the placental *hIGF1* nanoparticle gene therapy on cardiovascular function in offspring.

