## Supplemental Material for "Placenta *hIGF1* nanoparticle treatment in guinea pigs mitigates fetal sex dependent FGR-associated effects on kidney structure and blood pressure-related signaling pathways"

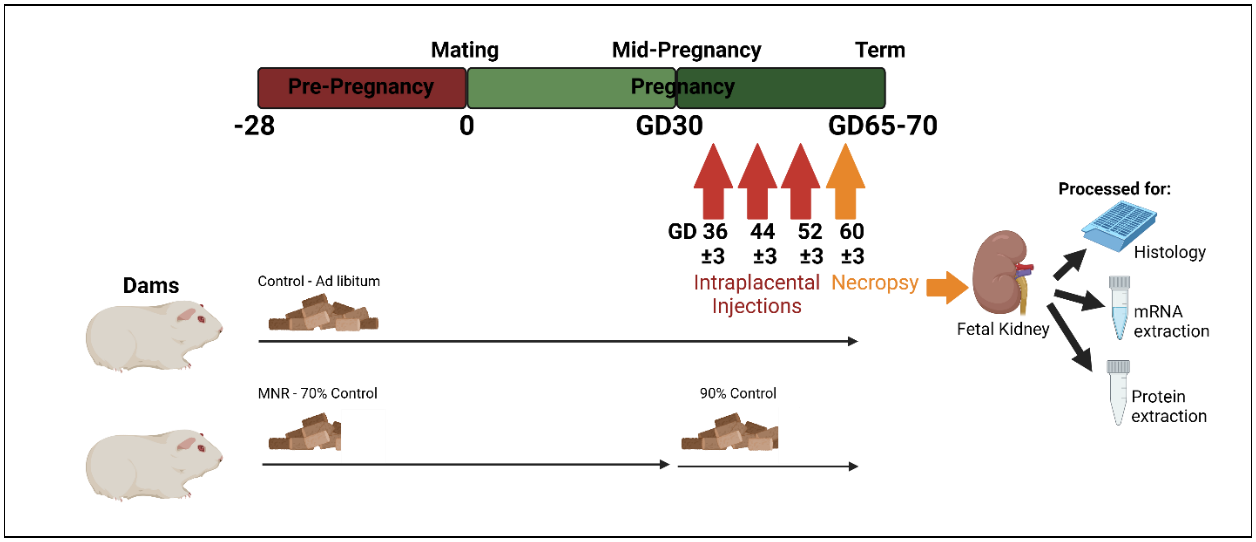


**Supplemental Diagram 1. Schematic of the experimental design showing Maternal Nutrient Restriction and Nanoparticle Treatment protocols.** Adapted from Davenport et al., 2025 [31]

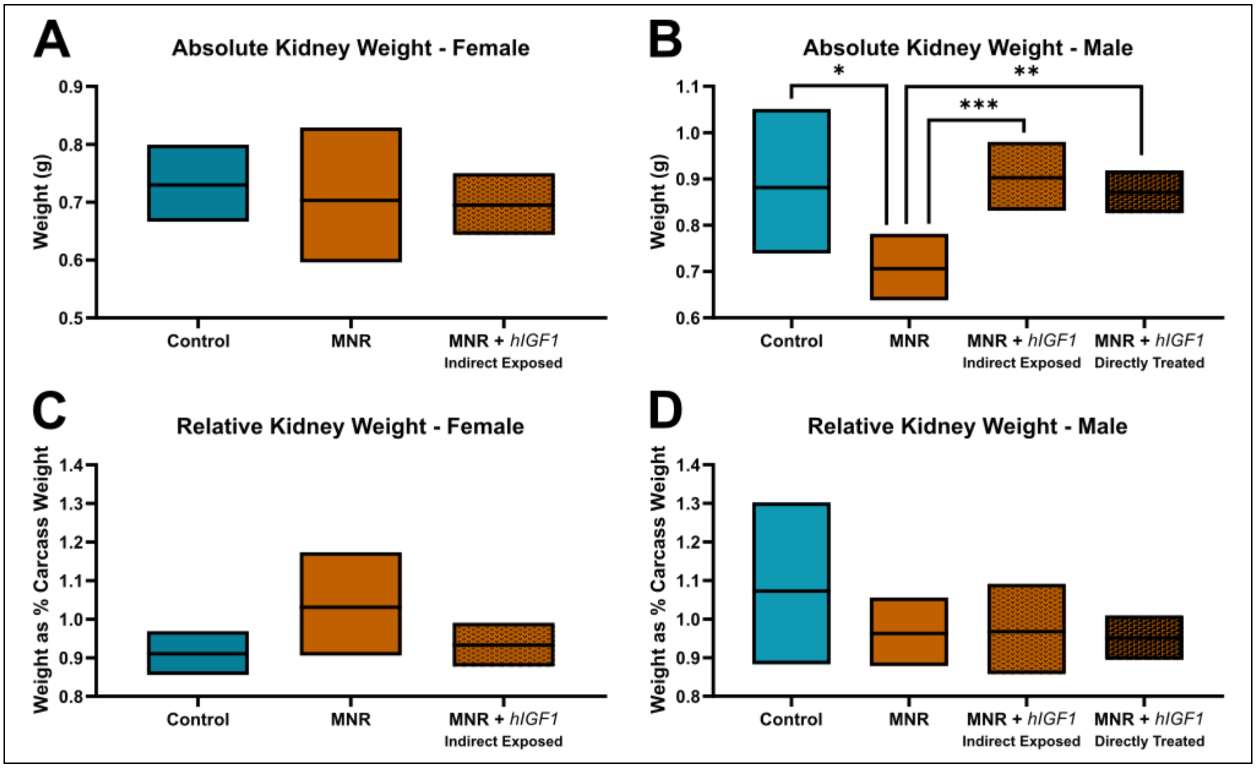


**Supplemental Figure S1. Effects of maternal nutrient restriction (MNR) and repeated placental *hIGF1* nanoparticle treatment (MNR + *hIGF1*) on fetal kidney weight. A**. In female fetuses, absolute kidney weight (both kidneys combined) was not different between sham nanoparticle Control, sham nanoparticle MNR and indirectly exposed MNR + *hIGF1* nanoparticle **B**. In female fetuses, kidney weight corrected for fetal weight was not different between sham nanoparticle Control, sham nanoparticle MNR and indirectly exposed MNR + *hIGF1* nanoparticle **C**. In male fetuses, absolute kidney weight (both kidneys combined) was reduced in the sham nanoparticle MNR compared to sham nanoparticle Control, and MNR + *hIGF1* nanoparticle **D**. In male fetuses, kidney weight corrected for fetal weight was not different between sham nanoparticle Control, sham nanoparticle MNR and MNR + *hIGF1* nanoparticle. Control: n = 6 dams (8 female and 11 male fetuses), MNR: n = 6 dams (5 female and 11 male fetuses), MNR + *hIGF1*: n = 5 dams (6 female and 10 male fetuses). Data are estimated marginal means ± 95% confidence interval. *P≤0.05; **P≤0.01. ***P≤0.001

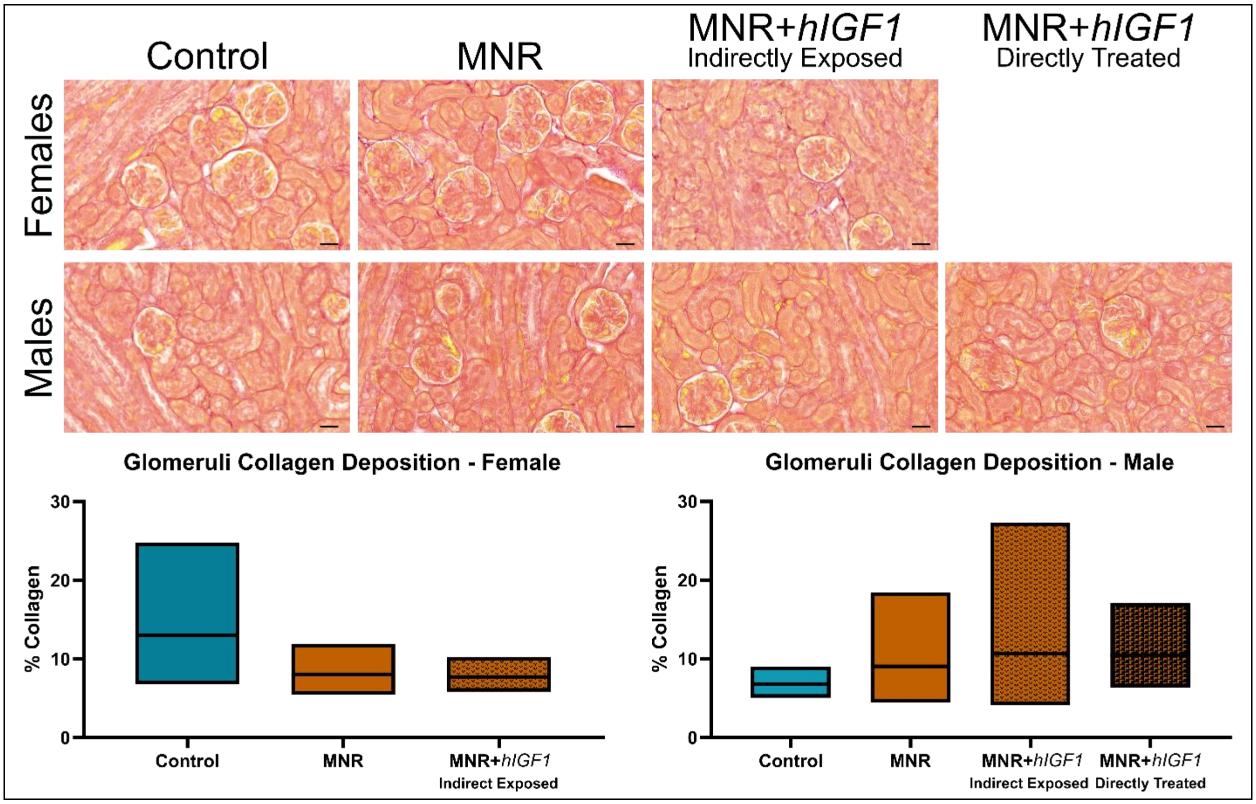


**Supplemental Figure S2. Representative images of collagen deposition (Sirius red stain) in the cortex and around the glomeruli of the near-term guinea pig kidney.** Quantification of the percentage collagen in the cortex and around the glomeruli indicated similar levels of collage between sham Control, sham maternal nutrient restriction (MNR) and MNR+*hIGF1* nanoparticle treated groups in both female male fetuses. 10 randomly selected fields of view were analyzed using ImageJ Software. Control: n = 6 dams (8 female and 11 male fetuses), MNR: n = 6 dams (5 female and 11 male fetuses), MNR+*hIGF1*: n = 5 dams (6 female and 10 male fetuses). Data are estimated marginal means ± 95% confidence interval. Scale bar = 50 µm

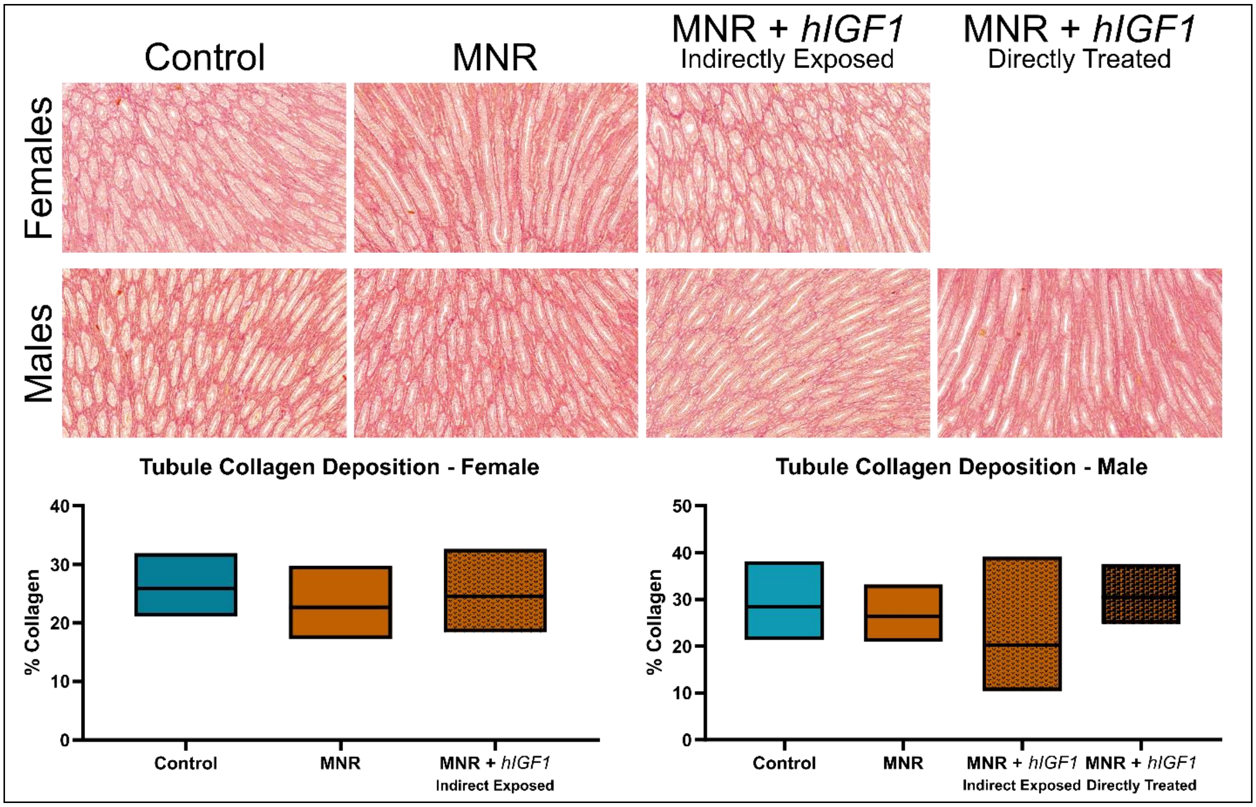


**Supplemental Figure S3. Representative images of collagen deposition (Sirius red stain) in the tubules of the near-term guinea pig kidney.** Quantification of the percentage collagen in the tubules indicated similar levels of collage between sham Control, sham maternal nutrient restriction (MNR) and MNR+*hIGF1* nanoparticle treated groups in both female male fetuses. 10 randomly selected fields of view were analyzed using ImageJ Software. Control: n = 6 dams (8 female and 11 male fetuses), MNR: n = 6 dams (5 female and 11 male fetuses), MNR+*hIGF1*: n = 5 dams (6 female and 10 male fetuses). Data are estimated marginal means ± 95% confidence interval. Scale bar = 50 μm

**Supplemental Table S1.** Fetal kidney expression of renin-angiotensin system, inflammation and oxidative stress-related genes and proteins

| | Control | | | MNR | | | MNR+*hIGF1* (Indirect Exposed) | | | MNR+*hIGF1* (Direct Inject) | | | P value | | |
|---|---|---|---|---|---|---|---|---|---|---|---|---|---|---|---|
| | | | | | | | | | | | | | Diet | Treat ment | Direct Injection (males only) |
| **Females** | | | | | | | | | | | | | | | |
| | EMM | 95% CI | | EMM | 95% CI | | EMM | 95% CI | | | | | | | |
| *Renin-Angiotensin System (Relative Quantification)* | | | | | | | | | | | | | | | |
| *RenBP* | 1.18 | 1.43 | 0.98 | 1.45 | 1.90 | 1.11 | 1.26 | 1.42 | 1.11 | | | | NS | NS | |
| *AgtR1* | 0.92 | 1.03 | 0.81 | 1.03 | 1.45 | 0.74 | 0.87 | 0.91 | 0.83 | | | | NS | NS | |
| *Inflammation and Oxidative Stress (Relative Quantification)* | | | | | | | | | | | | | | | |
| *Il6* | 3.31 | 1.91 | 5.74 | 1.31 | 0.59 | 2.90 | 2.40 | 1.58 | 3.64 | | | | NS | NS | |
| *Il6R* | 1.24 | 0.96 | 1.60 | 1.46 | 1.02 | 2.10 | 1.40 | 1.03 | 1.92 | | | | NS | NS | |
| *Hif1a* | 1.97 | 1.56 | 2.47 | 3.11 | 2.38 | 4.07 | 2.63 | 2.18 | 3.16 | | | | 0.022 | NS | |
| *Hspa5* | 2.57 | 1.93 | 3.42 | 6.74 | 4.71 | 9.62 | 4.02 | 3.06 | 5.28 | | | | <0.001 | 0.026 | |
| Catalase | 5.65E+05 | 3.25E+05 | 9.80E+05 | 3.94E+05 | 2.62E+05 | 5.93E+05 | 4.04E+05 | 2.12E+05 | 7.68E+05 | | | | NS | NS | |
| Thioreodoxin | 1.59E+06 | 6.88E+05 | 3.68E+06 | 1.49E+06 | 7.76E+05 | 2.85E+06 | 4.28E+05 | 2.41E+05 | 7.60E+05 | | | | NS | 0.002 | |
| SOD2 | 1.22E+05 | 8.82E+04 | 1.70E+05 | 1.89E+05 | 8.84E+04 | 4.04E+05 | 1.41E+05 | 5.33E+04 | 3.71E+05 | | | | NS | NS | |
| **Males** | | | | | | | | | | | | | | | |
| | EMM | 95% CI | | EMM | 95% CI | | EMM | 95% CI | | EMM | 95% CI | | | | |
| *Renin-Angiotensin System (Relative Quantification)* | | | | | | | | | | | | | | | |
| *RenBP* | 1.73 | 0.06 | 1.87 | 1.46 | 0.19 | 1.88 | 1.25 | 0.17 | 1.62 | 1.69 | 0.40 | 2.68 | NS | NS | NS |
| *AgtR1* | 0.95 | 1.55 | 0.58 | 0.81 | 0.93 | 0.70 | 0.98 | 1.55 | 0.61 | 0.56 | 0.68 | 0.45 | 0.026 | NS | NS |
| *Inflammation and Oxidative Stress (Relative Quantification)* | | | | | | | | | | | | | | | |
| *Il6* | 2.63 | 1.91 | 3.63 | 1.66 | 1.02 | 2.71 | 3.20 | 0.97 | 10.54 | 2.16 | 1.15 | 4.06 | NS | NS | NS |
| *Il6R* | 1.45 | 1.03 | 2.05 | 1.17 | 0.83 | 1.65 | 0.97 | 0.80 | 1.18 | 1.34 | 0.95 | 1.90 | NS | NS | 0.027 |
| *Hif1a* | 2.44 | 1.93 | 3.08 | 2.04 | 1.64 | 2.54 | 3.34 | 2.22 | 5.03 | 2.05 | 1.44 | 2.92 | NS | 0.027 | 0.014 |
| *Hspa5* | 4.23 | 3.29 | 5.44 | 3.85 | 3.03 | 4.88 | 5.38 | 4.11 | 7.04 | 4.64 | 2.51 | 8.57 | NS | NS | NS |
| Catalase | 5.06E+05 | 3.90E+05 | 6.56E+05 | 6.57E+05 | 4.37E+05 | 9.86E+05 | 2.74E+05 | 1.36E+05 | 5.53E+05 | 4.18E+05 | 3.44E+05 | 5.08E+05 | NS | 0.038 | NS |
| Thioreodoxin | 2.14E+06 | 1.45E+06 | 3.15E+06 | 2.20E+06 | 1.67E+06 | 2.89E+06 | 1.22E+06 | 4.75E+05 | 3.11E+06 | 1.19E+06 | 8.09E+05 | 1.75E+06 | NS | 0.045 | NS |
| SOD2 | 3.13E+05 | 1.80E+05 | 5.43E+05 | 1.19E+05 | 7.78E+04 | 1.82E+05 | 2.22E+05 | 6.63E+04 | 7.40E+05 | 9.74E+04 | 7.32E+04 | 1.30E+05 | 0.023 | NS | NS |

CI = Confidence Interval. EMM = Estimated Marginal Mean. MNR = Maternal Nutrient Restriction. Control and MNR are sham treated. Control: n = 6 dams (8 female and 11 male fetuses), MNR: n = 6 dams (5 female and 11 male fetuses), MNR+*hIGF1*: n = 5 dams (6 female and 10 male fetuses). Statistical significance calculated using Generalized Estimating Equations.
